# Song divergence and a gleaming white iris reveal four species in a widespread Neotropical understory bird (*Clibanornis rubiginosus*, Furnariidae)

**DOI:** 10.64898/2026.08.24.746896

**Authors:** Juan Carlos Villamizar, Andrés M. Cuervo

## Abstract

Polytypic species with large ranges may harbor unrecognized diversity because taxonomy ranks populations differing subtly in plumage as subspecies. The Ruddy Foliage-gleaner (*Clibanornis rubiginosus*) exemplifies this problem. It ranges from Mexico to Brazil, with 15 subspecies, and forms a non-monophyletic complex with two congeners, yet its songs had not been compared. We measured ten spectral and temporal variables on 104 recordings covering 14 of 15 subspecies. Bayesian linear mixed models showed three song groups: eight subspecies west of the Andes share a single-note song, whereas Amazonian and Guianan populations add a short introductory note and sing longer, lower-pitched songs. Within this group, *watkinsorum* sings the lowest-pitched and longest song and is phylogenetically closer to *C. cinnamomeigula* than to its Amazonian neighbors. The third group is *C. cinnamomeigula* alone, a white-eyed taxon in an otherwise dark-eyed group. Its high-pitched, vibrato song resembles none other in the genus. One-note and two-note songs differ in kind without intermediates, and every two-note taxon sequenced to date falls in one clade, so *C. rubiginosus* is paraphyletic. We recognize four species, *C. rubiginosus* sensu stricto, *C. cinnamomeigula*, *C. watkinsorum*, and *C. obscurus*. This raises *Clibanornis* from five species to eight and divides its only polytypic species.

## Introduction

“South America has been called quite rightly the ‘bird continent,’ as the number and variety of its feathered inhabitants surpass those of all other tropical land areas of the world.” (Haffer 1974, p. 51).

Much of this exceptional richness is the outcome of a long evolutionary history shaped by mountain building, the closure of isthmuses, and climatic cycles (Smith et al. 2014, Harvey et al. 2020). Uplift of the Andes, in particular, has long been recognized as an engine of diversification because it isolated populations, increased environmental gradients, and created new habitats (Chapman 1917, Fjeldså et al. 2012); Pleistocene climatic fluctuations later reinforced those effects (Haffer 1985, Flantua et al. 2019). Much of the resulting species diversity nevertheless remains underestimated, although largely described as subspecies, particularly in groups with subtle plumage variation (d’Horta et al. 2013, Dickens et al. 2021, Martínez-Gómez et al. 2023). In such groups, species limits are difficult to define from traditional plumage characters or external morphometrics, characters that are often evolutionarily conserved or that overlap extensively among populations (Krabbe and Schulenberg 1997, Lima and Vaz 2025).

This taxonomic challenge raises a basic question: how does divergence between species express itself when the visible phenotype offers few clues? Some birds differ markedly in plumage or morphology (Lara et al. 2012, Bocalini and Silveira 2016), whereas others are closely similar in appearance yet deeply divergent in acoustic or genetic signals (Chesser et al. 2020, Martínez-Gómez et al. 2023). Among widely distributed Neotropical birds, cryptic species diversity of this kind is often associated with physiographic features that fragment populations and act as barriers to gene flow (Smith et al. 2014). How much divergence accumulates across barriers such as the Andes or the large Amazonian rivers remains poorly documented (Harvey et al. 2020, Lees et al. 2020).

Acoustic signals are a key phenotypic character in species delimitation (Isler et al. 1998, Catchpole and Slater 2008), particularly in suboscine passerines. In that group, cultural learning of songs is limited, which makes them informative for inferring evolutionary relationships and species limits, especially when combined with molecular data (Remsen 2005, Isler et al. 2012). These vocalizations are directly involved in reproductive isolating mechanisms such as mate choice and territorial defense, and they can diverge during speciation (Areta and Pearman 2009, Fernández-Gómez et al. 2021) even where morphology does not (Martínez-Gómez et al. 2023).

This evidence has accumulated across several suboscine groups. Among antbirds, complexes whose members are morphologically exceedingly similar contain vocally diagnosable species (Isler et al. 1998, 2012, Chaves et al. 2010). The same pattern is documented in tapaculos (Krabbe and Schulenberg 2003, Cuervo et al. 2005), antpittas (e.g., Carantón-Ayala and Certuche-Cubillos 2010, Carneiro et al. 2019, Isler et al. 2020), tityrids (e.g., Musher et al. 2023, Lima et al. 2024), tyrant flycatchers (e.g., Cuervo et al. 2014, Lima and Vaz 2025, Bocalini et al. 2026), and ovenbirds (Zimmer 2002, Krabbe 2008, Cooper and Cuervo 2017). Song therefore often uncovers species diversity that had gone underestimated.

Across the Furnariidae, numerous taxa are taxonomically difficult because of their wide distributions and their close similarity in appearance and coloration, behavior and habitat use (Remsen 2003, Derryberry et al. 2011, d’Horta et al. 2013, Cooper and Cuervo 2017). For instance, molecular data showed that the foliage-gleaner genus *Automolus*, as traditionally defined, was not monophyletic: *A. rubiginosus* and *A. rufipectus* are more closely related to *Clibanornis* and *Hylocryptus* than to other *Automolus* (Derryberry et al. 2011, Claramunt et al. 2013). Both taxa were therefore transferred to *Clibanornis,* which now comprises five species of humid montane, premontane, and seasonal forests in the foothills of Central and South America, where they forage near the ground and in leaf litter (Remsen 2003).

Four out of five *Clibanornis* are monotypic, whereas *C. rubiginosus* shows the greatest geographic and phenotypic variation (Fig. 1). The most recent change in the complex separated *C. rufipectus* and raised it to species rank (Krabbe 2008) based vocal differences. Specifically, its songs resemble those of *C. erythrocephalus* more than those of any population of what was then *C. rubiginosus,* at the time still treated as an *Automolus.* Later phylogenetic work did not recover populations of *C. rubiginosus* as monophyletic with respect to *C. rufipectus* and *C. erythrocephalus* (Claramunt et al. 2013), a result since confirmed with genome-scale data (Harvey et al. 2020), which points to unrecognized diversity within the group. The number of recognized subspecies (15 sensu Remsen 2003), the range from Mexico south to Bolivia and east to French Guiana and northern Brazil (Fig. 2), and the absence of vocal and genetic data for several of them have so far prevented any comprehensive assessment of the complex.

**Figure 1.**
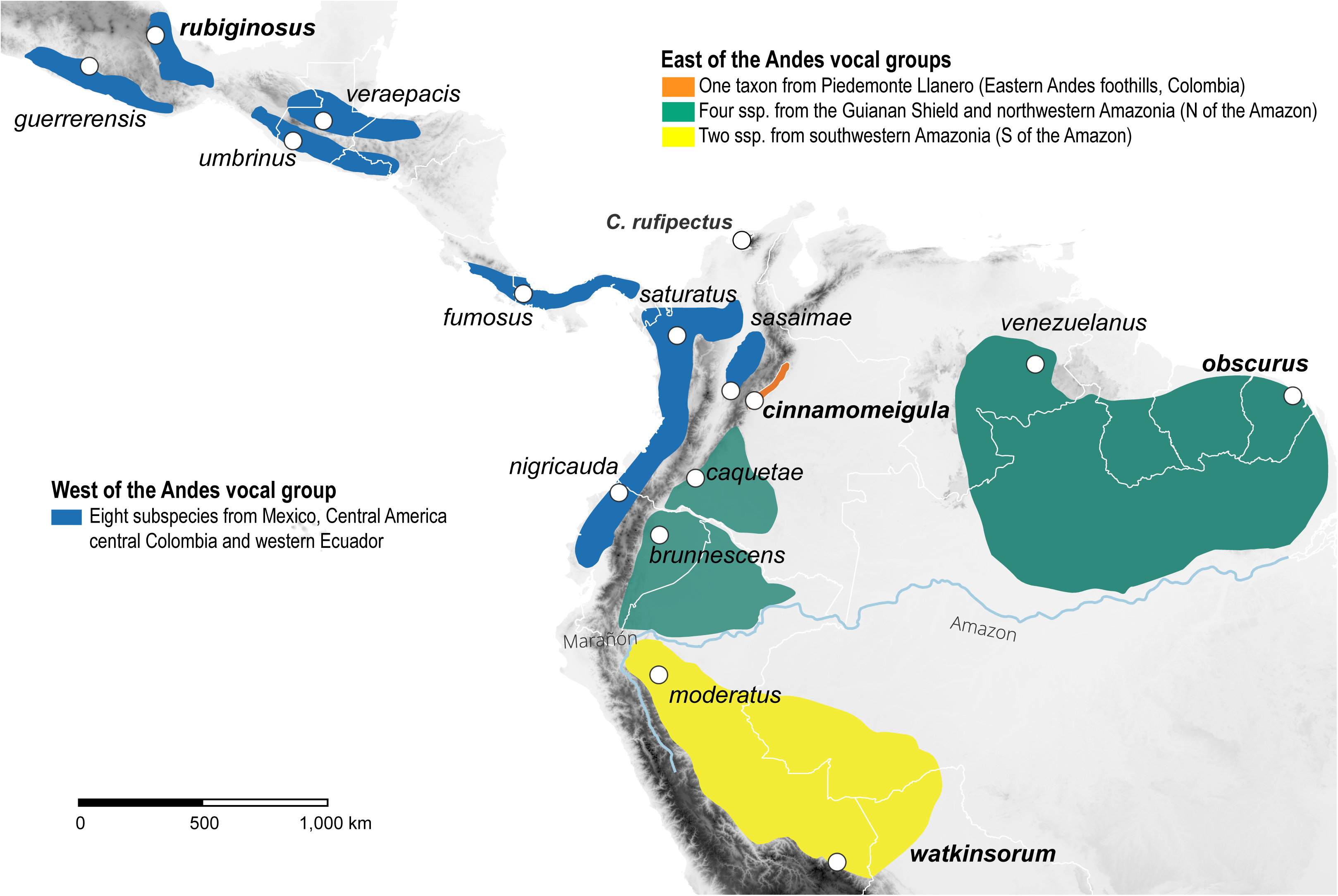
Geographic distribution of the Ruddy Foliage-gleaner (*Clibanornis rubiginosus*) and Santa Marta Foliage-gleaner (*C. rufipectus*). Polygons show the approximate range in the complex of each recognized taxon and grouped according to this study. Fill color shades denote the approximate range of subspecies vocal groups: blue (Mexico through Central America, Colombian inter-Andean valleys and south to Choco in Ecuador), orange (Llanos foothills at the base of the Eastern Andes, Colombia), teal (Guianan shield and northwestern Amazonia), yellow (western Amazonia, south of the Amazon). Circles mark the type locality of each subspecies, and of *C. rufipectus*.

**Figure 2.**
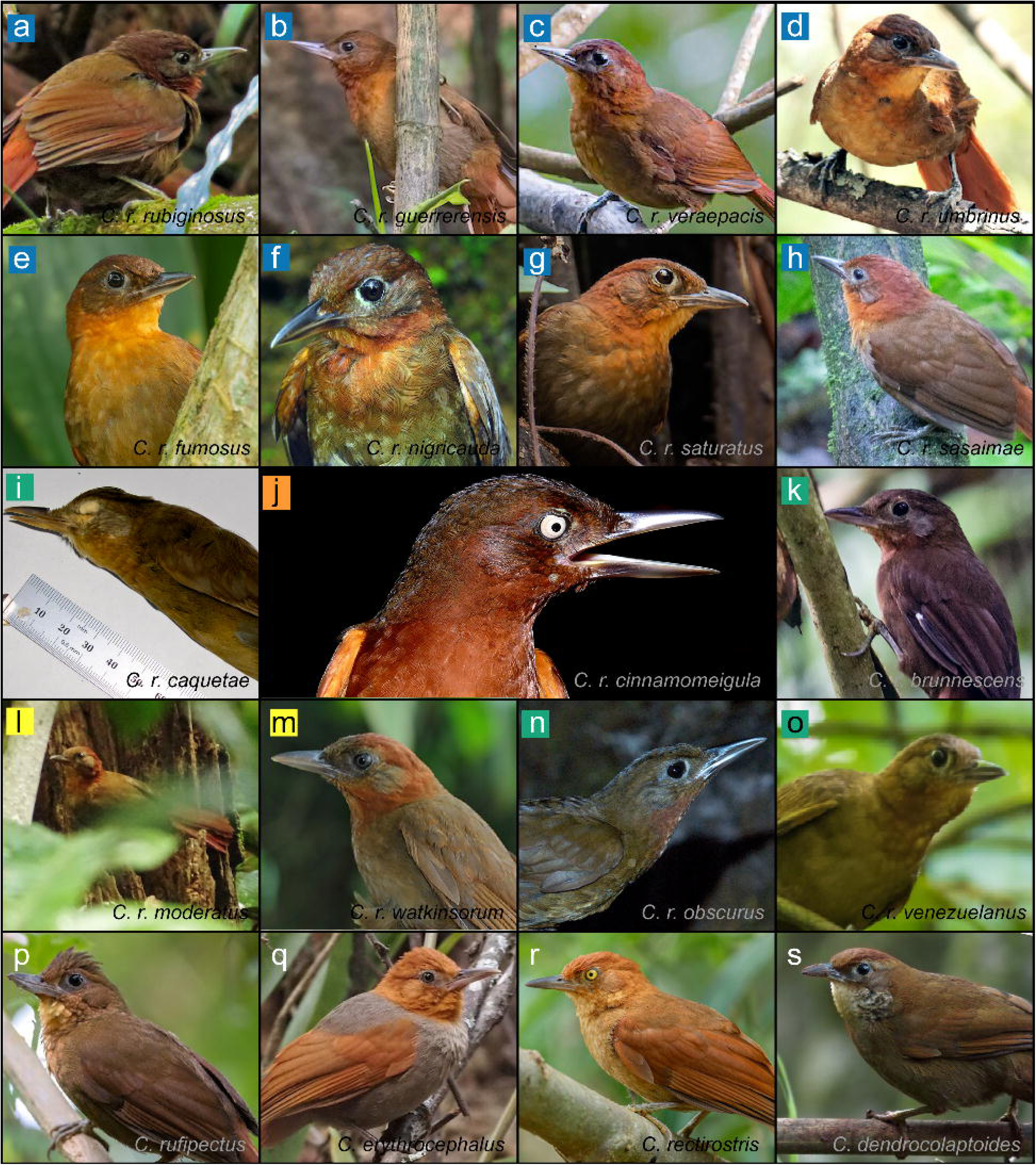
Photographs of each subspecies of the *Clibanornis rubiginosus* complex and of the four remaining species of *Clibanornis*. (a) *C. r. rubiginosus*, (b) *C. r. guerrerensis*, (c) *C. r. veraepacis*, (d) *C. r. umbrinus*, (e) *C. r. fumosus*, (f) *C. r. nigricauda*, (g) *C. r. saturatus*, (h) *C. r. sasaimae*, (i) *C. r. caquetae*, (j) *C. r. cinnamomeigula*, (k) *C. r. brunnescens*, (l) *C. r. moderatus*, (m) *C. r. watkinsorum*, (n) *C. r. obscurus*, (o) *C. r. venezuelanus*, (p) *C. rufipectus*, (q) *C. erythrocephalus*, (r) *C. rectirostris*, (s) *C. dendrocolaptoides*. Colored marks as in Fig. 1. Photographers are listed in the Supporting material.

Some subspecies described within *C. rubiginosus* vary in coloration, distribution, and size, which suggests heterogeneous degrees of evolutionary differentiation. The most striking case are that of *C. r. cinnamomeigula* of the foothills above the Llanos of Colombia, and *C. r. watkinsorum* of southwestern Amazonia (Fig. 2). These are two of the most intensely and distinctively colored members of the complex. The rufous cinnamon plumage that contrasts sharply with its pale, almost white irides in *C. r. cinnamomeigula* differs sharply from the dark brown irides of all other subspecies (Fig. 1j). This character, now one of the most diagnostic features of the taxon, went unnoticed for decades. The type is a Bogotá skin without field data (Hellmayr 1905), and specimens collected before 2005 carry no iris color on their labels. Only field observations and specimens taken from 2005 onward revealed the contrast between the pale iris, the dark face, and the deep cinnamon body (F. G. Stiles, A. M. Cuervo, J. P. López and O. Laverde, pers. obs.). On the other hand, *C. r. watkinsorum* is characterized by bright rufous nape and sides of head that separates them from the rest of the Amazonian populations (Fig. 1m).

In contrast, the other northwestern Amazonian and Guianan shield taxa are harder to distinguish. North of the Amazon river in the Peruvian and Ecuadorian Amazon, *C. r. brunnescens* (Fig. 1k) is a dark form nearly indistinguishable from *C. r. caquetae* of Colombia, although the latter is described to have a more rufous throat. Both *C. r. brunnescens* and *C. r. obscurus* of the Guianan Shield are smaller in size than subspecies south of the Amazon (e.g., watkinsorum, Hellmayr 1925, Meyer de Schauensee 1950). Taxonomic treatment of these populations has varied. *Clibanornis r. obscurus* and *C. r. watkinsi* have been treated as a distinct species with no rationale by the Brazilian Ornithological Records Committee (Piacentini et al. 2015), although global checklists have not adopted that treatment and retain it as a subspecies. West of the Andes the pattern is different. *Clibanornis r. nigricauda*, *C. r. saturata,* and *C*.

*r. sasaimae* (Fig. 1f-h), restricted to the foothills of the Chocó region and to the inter-Andean valleys of central Colombia, are the darkest taxa with and more contrasting chestnut colors than other members of the complex; they have been treated as distinct forms since Hellmayr (1925). In Central America and Mexico, the subspecies *rubiginosus, umbrinus, veraepacis, guerrerensis,* and *fumosus* vary more subtly in coloration, probably clinally (Salvin and Godman 1891). This phenotypic heterogeneity, together with inconsistent taxonomic treatments and the absence of bioacoustic analyses, suggests that *C. rubiginosus* sensu lato represents a complex of lineages at different stages of differentiation whose limits require an integrative assessment.

Recent taxonomic progress, including the redefinition of *Automolus* and *Clibanornis* (Claramunt et al. 2013) and the treatment of *C. rufipectus* at species rank on vocal evidence (Krabbe 2008), left the status of populations within the *C. rubiginosus* complex unresolved. Missing recordings for most subspecies and limited genetic sampling have prevented a rigorous delimitation, and no previous study has compared songs across the complex. We analyze all recognized subspecies except *C. r. caquetae* in order to (1) characterize geographic vocal variation across the species range and identify vocally diagnosable groups, and (2) evaluate whether those groups correspond to current taxonomic limits or point to unrecognized species-level taxa. We document vocal divergence structured largely by the Andes, with two further groups east of the cordillera, which raises the number of species in *Clibanornis* from five to eight.

## Material and methods

### Recording selection and processing

We analyzed 104 recordings (Table S1) downloaded from the Macaulay Library (ML, Cornell Lab of Ornithology, Ithaca, NY, USA) and xeno-canto (XC, https://xeno-canto.org/). We focused on recordings of quality A (loud, clear) and B (relatively clear, little interference) and discarded quality C recordings (short, with excessive background noise, interference from other vocalizations, or adverse weather such as rain). To increase sample size, we included recordings obtained with playback. Because playback is used frequently and because the databases often lack precise information about it, we selected, whenever possible, segments in which the song showed no sign of agitation (e.g. songs of normal intensity, without marked acceleration of delivery rate). We assumed that recordings from the same date and locality corresponded to the same individual.

Sample size varied among subspecies. We compiled ten recordings each for *brunnescens, fumosus, nigricauda, obscurus, saturatus, umbrinus,* and *watkinsorum;* seven each for *guerrerensis* and *rubiginosus;* six each for *cinnamomeigula, sasaimae,* and *veraepacis;* and one each for *moderatus* and *venezuelanus.* No confirmed recordings exist for *caquetae,* a subspecies of the Amazonian foothills of southern Colombia (Meyer de Schauensee 1947). The sexes of *C. rubiginosus* are alike in plumage, and the sex of recorded birds is not known, so we could not assign sex to any recording.

We normalized the recordings and resampled them to a single mono channel at 44.1 kHz and 16 bits in WAVE format using Audacity 3.6.3 (Audacity Team 2024). We measured four song bouts per recording to capture within-individual variation, defining about as one complete song delivered once, and analyzed them independently in the quantitative analyses. We took measurements manually from spectrograms in Raven Pro 1.6 (K. Lisa Yang Center for Conservation Bioacoustics 2024) using a 512-sample Hann window, a 3 dB filter bandwidth of 124 Hz, and 50% overlap. We adjusted brightness (50–70%) and contrast individually for each recording to optimize spectrogram display.

### Acoustic variables

We defined the song of *C. rubiginosus* as a simple nasal sound 0.3–0.8 s long, composed of one or two notes. Two-note songs comprise a short note (< 0.2 s) followed by a longer main note, separated by a brief silence (0.01–0.1 s). Song bouts typically consist of one to three repetitions of this basic structure (Fig. 3). To capture the full vocal variation of the group, we measured ten spectral and temporal variables, both for the complete song and for its two main structural components (subunit 1 and subunit 2). We defined a note as a continuous trace of acoustic energy on the sonogram (Catchpole and Slater 2008). In two-note songs, we measured the introductory note (subunit 1) and the main note (subunit 2) separately. In single-note songs we took all measurements on the entire note. The acoustic parameters measured were (1) total duration, (2) 90% duration, (3) maximum time, (4) center time, (5) frequency range (delta frequency), (6) 5% frequency, (7) 95% frequency, (8) peak frequency (frequency of maximum amplitude), (9) 90% bandwidth, and (10) center frequency. We chose the 5% and 95% frequencies rather than minimum and maximum frequencies because they are more clearly defined on the spectrogram and less sensitive to background noise. We generated the most representative spectrograms, from the highest-quality recordings (Figs 2 and 3), with the R package seewave (Sueur et al. 2008).

**Figure 3.**
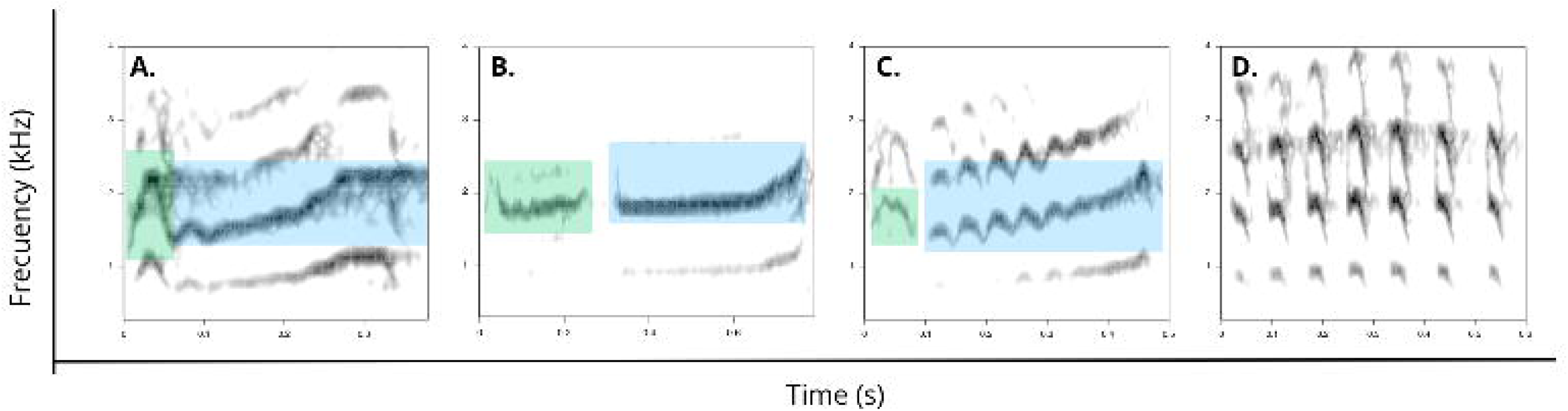
Representative spectrograms of the song of *Clibanornis rubiginosus*. (a) Western group (*C. r. fumosus*, Puntarenas, Costa Rica, ML39298). (b) Eastern group in western Amazonia (*C. r. watkinsorum*, Madre de Dios, Peru, ML76198). (c) *Clibanornis r. cinnamomeigula* from the piedemonte llanero (Boyacá, Colombia, ML126777531). (d) *Clibanornis rufipectus* from the Sierra Nevada de Santa Marta (XC101808). Green boxes delimit subunit 1 (introductory note), blue boxes subunit 2 (main note); the box outline is heavier on the subunit of greatest energy. Panels (a) to (c) show one complete song of each group.

To assess the sampling adequacy of the correlation matrix, we computed the Kaiser-Meyer-Olkin (KMO) measure using the R package *psych* (Revelle 2026), and then we inspected the correlation matrix visually with corrplot (Wei and Simko 2026). We retained only those variables with an individual measure of sampling adequacy (MSA) ≥ 0.6, until reaching an overall KMO ≥ 0.7 (Budaev 2010, Fernández-Gómez et al. 2021). This procedure removed five variables (maximum time, center time, 90% duration, delta frequency, and 90% bandwidth) and retained the remaining five. We then used a principal component analysis (PCA) with the *prcomp* function in R to reduce the dimensionality of the data (Budaev 2010).

### Statistical analysis

We analyzed vocal variation within the *C. rubiginosus* complex between the one-note and two-note groups, and among the 14 sampled subspecies (all except *caquetae),* with Bayesian linear mixed models (BLMM) fitted by Hamiltonian Monte Carlo and the no-U-turn sampler (NUTS) in the package *brms* (Bürkner 2017) in R 4.4.2 (R Core Team 2024). We chose Bayesian mixed models because they handle unequal sample sizes and accommodate random effects flexibly (Hadfield 2010). We used a Gaussian error distribution and the default priors of brms. We report 95% highest posterior density intervals (HDI), and we treated an interval that excludes zero as evidence for a difference.

To evaluate vocal variation between groups (one note vs two notes), we fitted BLMMs with five different response variables at three levels of analysis (complete song, subunit 1, and subunit 2), for a total of 15 models (Table S2). We modeled the five vocal traits as response variables: total duration, center frequency, peak frequency, 5% frequency, and 95% frequency. In all models we treated the number of notes as a fixed effect and the identification code of each recording as a random effect, to control for pseudoreplication arising from the four measurements per individual (Bolker et al. 2009). We standardized all response variables to a Z scale with the scale function of the base package, which gave a mean of 0 and a standard deviation of 1 and allowed us to compare effect sizes among models (Nakagawa and Cuthill 2007). We also used the two components extracted by principal component analysis, PC1 (spectral) and PC2 (temporal), to explore vocal diversity in the group.

We ran all analyses for 50,000 iterations with a 20% warm-up and thinning every 20 iterations to reduce autocorrelation between consecutive samples, which yielded 8000 posterior samples per model (4 chains × 2000 samples). We checked chain convergence with the R-hat statistic and calculated the effective sample size (ESS); all parameters had ESS > 3000 and R-hat < 1.01 (Table S3). We then made pairwise comparisons at the group and subspecies levels with the contrast function in the package emmeans (Lenth and Piaskowski 2026), calculating posterior differences and their 95% HDI.

## Results

We identified three dominant song types in the *C. rubiginosus* complex that correspond to three well-differentiated geographic groups (Fig. 3). The Eastern Cordillera of the Andes is the main barrier separating two of these groups, and it harbors the third (Fig. 4). The western group (n = 8 subspecies, 66 recordings) ranges from southern Mexico east to the southern end of the Chocó in western Ecuador and through the inter-Andean valleys to central Colombia. This group, comprising the subspecies *rubiginosus, guerrerensis, veraepacis, umbrinus, fumosus, saturatus, sasaimae,* and *nigricauda,* gives a song of a single note shaped like an elongated inverted V that rises at the end. Total song duration is 0.43 ± 0.13 s (mean ± SD), with frequencies from 1598 ± 260 Hz at the start to 2087 ± 187 Hz at the peak. The note is repeated one to three times; when two or three repetitions follow one another, intervals between notes range from 0.2 to 0.6 s, whereas in isolated deliveries the intervals reach 1.2–2.5 s.

**Figure 4.**
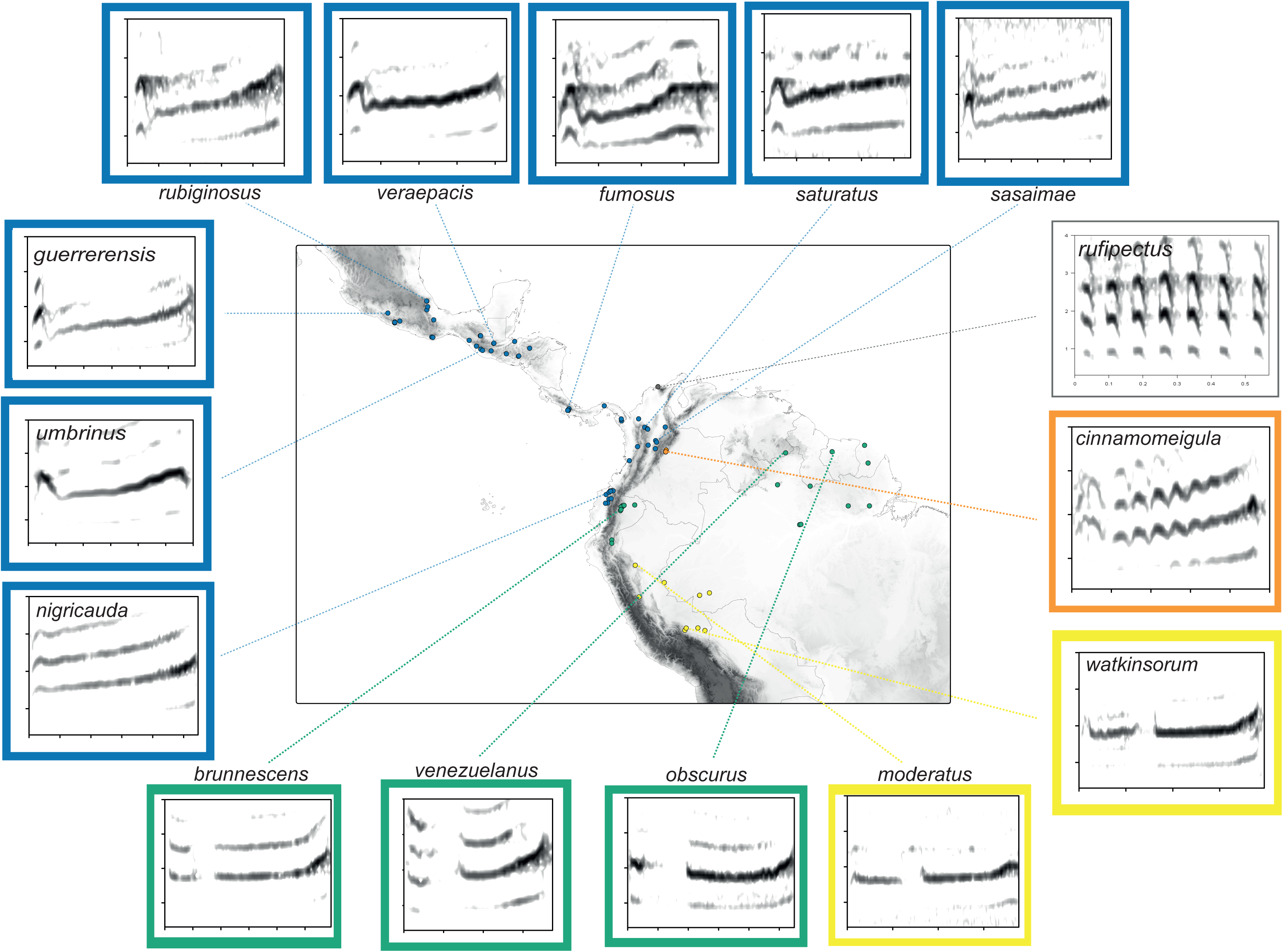
Geographic variation in song across the *Clibanornis rubiginosus* complex. Each taxon is represented by one spectrogram, except *caquetae*, for which no recording is available. Axes are common to all panels. Lines connect each spectrogram to its geographic origin. Outline colors denote vocal group, and as in previous figures: blue, western group (*C. rubiginosus* sensu stricto); orange (*C. cinnamomeigula*); yellow, southwestern Amazonia group (*C*. watkinsorum, including *moderatus*); teal, eastern group in northwestern Amazonia and Guiana shield (*C. obscurus*); grey, *C. rufipectus*.

The eastern group (n = 5 subspecies, 32 recordings) comprises the subspecies *brunnescens, venezuelanus, obscurus, watkinsorum* and *moderatus,* distributed east of the Andes in the Amazonian foothills, the Amazonian lowlands, and the Guianan Shield. These populations show a vocal pattern different from that of the western group, with two-note songs. The song begins with a very short introductory note (0.13 ± 0.06 s), followed by a silent interval (0.03–0.09 s), and ends with a longer second note (0.35 ± 0.07 s) that rises slightly at the end. Total song duration is 0.55 ± 0.11 s, with frequencies from 1616 ± 186 Hz at the start to 1953 ± 206 Hz at the peak; peak and center frequencies are therefore lower than in the western group. Within this vocal structure, the songs of *watkinsorum* and *moderatus* are diagnosable from those of *brunnescens, venezuelanus and obscurus* (see below).

The *cinnamomeigula* group (n = 1 subspecies, 6 recordings) is a third, strongly divergent vocal group restricted to the foothills above the Llanos on the eastern versant of the Eastern Cordillera of Colombia. Although it shares a two-note structure with the eastern group, its components differ structurally from those of any other taxon in the complex. The introductory note (subunit 1) is a brief, dry sound of less pure tone with a clicking quality (0.07 ± 0.02 s), separated from the main note (subunit 2) by a very short silence (0.01–0.02 s). The main note shows a distinctive frequency modulation organized in series of five to seven rapid serrated oscillations that produce a resonant vibrato (0.37 ± 0.05 s). The complete song lasts 0.48 ± 0.05 s and reaches the highest peak frequency of the entire complex (2139 ± 243 Hz).

The principal component analysis showed that the first component (PC1), dominated by frequency variables, explained 41% of the total acoustic variance and structured the differentiation among groups mainly along a frequency axis. The second component (PC2) explained 18.4% of the variance and was associated mainly with temporal variables, including song duration. This ordination was exploratory; statistical inference about differences among groups rests on the Bayesian linear mixed models, which we ran on the original acoustic variables.

The Bayesian linear mixed models comparing the one-note (western) and two-note (eastern) groups differed in spectral and temporal variables (Fig. 5, Table S3). The two-note group had lower peak frequency (PF) and center frequency (CF) than the one-note group, with 95% HDIs that exclude zero (PF: estimate = -0.70, 95% HDI -1.08 to -0.33; CF: estimate = -0.66, 95% HDI -1.04 to -0.28). Total song duration (TD) was greater in the two-note group (estimate = 0.97, 95% HDI 0.58–1.35), again with an HDI that excludes zero. We detected no difference in 5% frequency (estimate = 0.07, 95% HDI -0.33 to 0.47). The 95% frequency tended toward lower values in the two-note group (estimate = -0.36, 95% HDI -0.75 to 0.03), but the HDI includes zero and we do not treat it as a difference.

**Figure 5.**
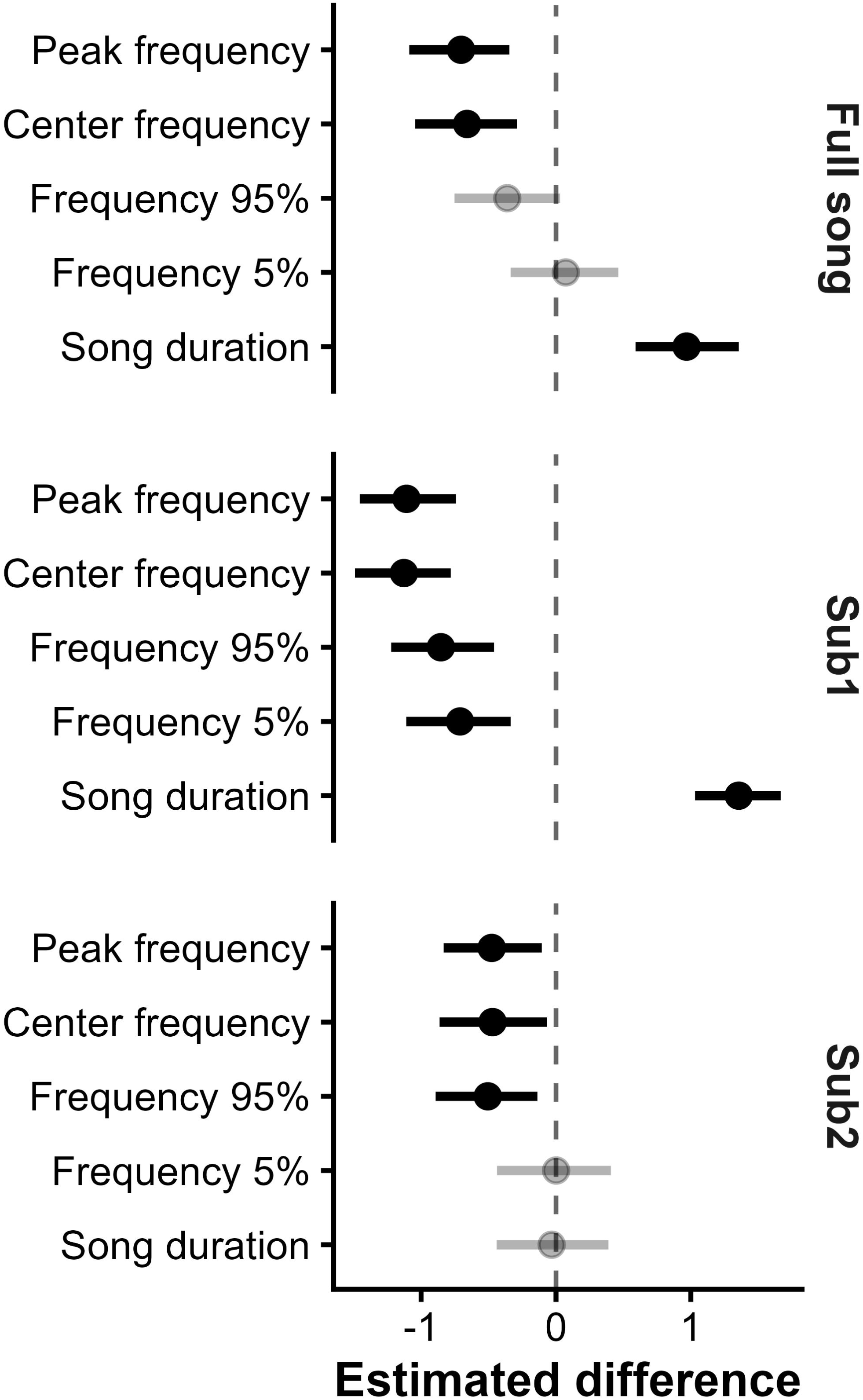
Differences in song traits between the two groups of *Clibanornis rubiginosus* separated by the Andes, shown as 95% highest posterior density intervals. Values are standardized differences between the two-note (eastern) and one-note (western) groups; positive values indicate higher values in the eastern group. Results are shown for the complete song and for subunits 1 and 2. *Clibanornis r. cinnamomeigula* was excluded from this analysis.

We excluded *C. r. cinnamomeigula* from the Bayesian model comparisons because its song type is unique and does not correspond to either the western or the eastern pattern. Its acoustic values differ in frequency from both main groups, with a peak frequency of 2139 ± 243 Hz, the highest recorded for the complex. Total duration of the *cinnamomeigula* song (0.48 ± 0.05 s) falls within the range of both groups. Within the western group, pairwise comparisons among the eight subspecies recovered no differences in any acoustic variable (all HDIs included zero) across a distribution spanning roughly 4500 km. The eastern group was more heterogeneous: *C*.

*r. watkinsorum* differed from the remaining eastern subspecies in its lower frequencies (peak frequency = 1838 ± 132 Hz), in the longest total song duration of the complex (0.66 ± 0.05 s), and in a longer introductory note (0.21 ± 0.02 s). The song of *moderatus* (n=1) was closest to *watkinsorum*, and all the remaining eastern subspecies (*brunnescens, obscurus, venezuelanus*) were acoustically indistinguishable from one another (all HDIs included zero).

## Discussion

Song structure within the *C. rubiginosus* complex fall into three vocal groups whose boundaries are largely defined by the Andes and their foothills. Eight subspecies west of the cordillera share a single-note song that varies subtly from Mexico to western Ecuador, despite spanning 4500 km. East of the Andes, populations add a short introductory note, but with two subspecies showing a further divergence. The western Amazonian *watkinsorum* (including *moderatus*) sings lower and longer than any other taxon in the complex, and its introductory note alone separates it from other Amazonian or Guianan shield population. The third group holds a single taxon, *cinnamomeigula*, restricted to the Llanos foothills of Colombia, which delivers a modulated, serrated vibrato found nowhere else in the genus. Song has therefore diverged along four species-level trajectories beneath a plumage color pattern that has changed comparatively little in what is currently treated as a single, polytypic species.

The character that divides the two main groups is categorical rather than a shift in means. Songs west of the Andes have one note, east of the Andes two, with an introductory note followed by a main note, and all 98 recordings assigned to these groups fall cleanly on one side of that boundary. Discrete vocal characters tend to carry more weight in delimitation than continuous variables, which can grade clinally without marking any limit (Isler et al. 1998, Patten and Unitt 2002). That the boundary coincides with the cordillera fits the role long attributed to the Andes as an engine of avian diversification in lowland birds, particularly for understory birds with limited dispersal (Chapman 1917, Fjeldså et al. 2012, Smith et al. 2014, Cadena et al. 2016).

The four groups are not just acoustic clusters, but each also corresponds to genetically defined evolutionary lineages: the Llanos foothill *cinnamomeigula* and the western Amazonian *watkinsorum* form a well-supported clade that stands apart from the trans-Andean subspecies and the other the Guianan Shield and remaining Amazonian populations (Claramut et al. 2013).

Harvey et al. (2020), using ultraconserved elements, also found that *cinnamomeigula* belongs with the populations east of the Andes and with none of those to the west. Therefore, all taxa east of the sings a two-note songs, so the introductory note is a shared state by descent. Under the current arrangement *C. rubiginosus* is not monophyletic with respect to either *C. rufipectus* or *C. erythrocephalus*, both nested among its subspecies (Claramunt et al. 2013, Harvey et al. 2020). Vocal evidence removed *rufipectus* from the complex (Krabbe 2008) without resolving vocal variation in the rest.

The two cross Andes groups are not equally variable within. Songs across the western group overlap strongly among subspecies in both temporal and spectral variables, and the absence of discrete clusters is more compatible with clinal variation than with structure, consistent with a preliminary assessment of plumage in which the Central American subspecies form a coherent, if not homogeneous, morphological group (Remsen 2003, Claramunt et al. 2013). One pattern in the ordination suggest that *C. r. fumosus* of Panama and Costa Rica is vocally closer to the South American populations (*saturatus*, *nigricauda*, and *sasaimae*) than to those of Mexico and northern Central America (*rubiginosus*, *guerrerensis*, *veraepacis*, and *umbrinus*), although the models recover no diagnostic difference among any of them. The pattern conflicts with that for plumage, which places *fumosus* with the nominate group (Claramunt et al. 2013). Comparable asymmetric affinities between lower Central America and South America are documented in other birds (Smith et al. 2014, Dickens et al. 2021, Moncrieff et al. 2022, Reyes et al. 2023). Because *C. rufipectus* and *C. erythrocephalus* fall near or within *C. rubiginosus* sensu stricto as defined herein, it may prove to be paraphyletic (Claramunt et al. 2013, Harvey et al. 2020). Our vocal data give no basis to subdivide it. Resolving its internal structure will require sampling across the isthmus and both Andean slopes, together with *C. rufipectus* and *C. erythrocephalus*.

The eastern group of Amazonia and the Guianan Shield, in contrast, is vocally heterogeneous, and *C. r. watkinsorum* accounts for most of that variation. It gives the lowest-pitched and longest song of the entire complex, not only of the eastern group, and its introductory note is longer and lower pitched than in any other eastern population, separating it from *brunnescens* and *obscurus* in a discriminant function analysis. The single recording available for *moderatus* falls with *watkinsorum* on both duration and peak frequency, although its sample size kept it out of that analysis. Zimmer (1935) interpreted *moderatus* as morphologically intermediate among several taxa of the complex rather than as an ally of *watkinsorum*, but later authors questioned its validity and treated it as part of a variable *watkinsi* (Remsen 2003), and the acoustic evidence points the same way. One recording, of course, cannot distinguish vocal affinity from individual variation, yet in our opinion the burden of proof now falls on retaining *moderatus* rather than on synonymizing it. Its phylogenetic position matches that acoustic distinctiveness. *Clibanornis watkinsorum* is sister to *C. cinnamomeigula* rather than to other Amazonian populations (Claramunt et al. 2013).

No population in the complex diverges as sharply as *C. r. cinnamomeigula*. Its song is unlike any other in the group, and this acoustic distinctiveness matches its genetic position. *Clibanornis cinnamomeigula* falls within a well-supported clade of taxa east of the Andes in both molecular studies available, with *watkinsorum* in Claramunt et al. (2013). Consistent vocal divergence of a comparable magnitude was sufficient to restore species rank to *C. rufipectus* (Krabbe 2008), and the same standard applied here points to an independent evolutionary lineage restricted to the Andean foothills above the Orinoco plains of Colombia. Its distinctiveness was recognized early; it was described as a species (Hellmayr 1905) and retained as such by Chapman (1917, 1926); it was subsumed into *rubiginosus* by Cory and Hellmayr (1925) and has been treated as a subspecies of broadly defined *rubiginosus* ever since. Its distinct iris coloration was not known until 2004–2007, when modern specimens where collected.

The song of *cinnamomeigula* also places it closer to other members of the genus than to *C. rubiginosus*. An introductory note followed by a main note modulated as a vibrato somewhat resembles the structure *C. rufipectus* and *C. erythrocephalus*. None of the three is the closest relative of the others (Claramunt et al. 2013, Harvey et al. 2020), so the shared structure cannot be interpreted as recent common ancestry. However, in *cinnamomeigula* the main note is delivered as a continuous unit with rapid frequency oscillations, whereas in *rufipectus* and *erythrocephalus* the song is articulated into discrete, separated notes. Continuous vibrato and pulsed notes may be two states of the same character, with *cinnamomeigula* exhibiting an intermediate position.

Four species should be recognized in what is currently treated as *C. rubiginosus*, a conclusion that follows the framework of de Queiroz (1998, 2007), in which species correspond to independent evolutionary lineages. The acoustic results demonstrate diagnostic advertising signals that reflect that independence under several of the criteria ornithologists apply. Songs are genetically inherited in suboscines, so consistent vocal diagnosability identifies distinct lineages. The same diagnosability bears on the biological species concept (Mayr 1963), because song functions in mate choice and territorial defense and can therefore restrict gene flow, although no contact zone between *cinnamomeigula*, *watkinsorum*, and the core populations has been documented and thus reproductive isolation cannot be tested directly. Phenotype reinforces the acoustic evidence. *Clibanornis cinnamomeigula* is uniformly saturated cinnamon below with a dark face and striking whitish-yellow irides, a character absent from every other taxon in the complex, all of which have dark brown irides (Fig. 1); *C. watkinsorum* has little rufous overall, dark scaling on the throat, and a rufous nape (Remsen 2003, Claramunt et al. 2013). Lineages diagnosable in song and plumage and independent in phylogeny are unambiguously recognized as species (Coyne and Orr 2004, de Queiroz 2007, Winker 2021).

*Clibanornis rubiginosus* sensu stricto (Sclater, 1857) then is formed by populations west of the Andes, the nominate subspecies together with *fumosus*, *guerrerensis*, *nigricauda*, *sasaimae*, *saturatus*, *umbrinus*, and *veraepacis*. The eastern assemblage takes the oldest name available, *Clibanornis obscurus* (Pelzeln, 1859), and includes all populations north of the Amazon river: *brunnescens*, *venezuelanus*, and, provisionally, *caquetae*. Neither proposed species is without precedent. Species rank for *obscurus*, with *venezuelanus* as a subspecies, and for *watkinsorum* is already in use for the Brazilian avifauna (Piacentini et al. 2015), and a preliminary assessment of plumage found no diagnostic differences among the four taxa we place in *C. obscurus* (Claramunt et al. 2013). Single recordings represent *venezuelanus* and *moderatus*, so the internal homogeneity we report for the eastern group rests mostly on *brunnescens* and *obscurus*, and no recording of *caquetae* exists, which leaves its placement uncertain.

Recognizing these lineages as species has conservation implications. *Clibanornis cinnamomeigula* is the critical case. It is confined to the Andean foothills above the Colombian Llanos, a narrow linear band of roughly 7000–8000 km² where agricultural expansion has driven some of the highest rates of land-cover change in the country (Sanchez-Cuervo et al. 2013, Briceño-Vanegas and Gallego-Herrera, 2025). A range that small under that much pressure makes the species likely threatened. Large gaps in the record across central Amazonia leave the range limits uncertain between *watkinsorum* and *obscurus,* although separated by the Amazon, and between *caquetae* and *cinnamomeigula*, and no contact zone has been sampled or identified. Acoustic sampling across subspecies, high-resolution genomic data, and material from the Andean-Amazonian transition where *caquetae* occurs would help resolve what remains open in this complex.

## Conclusion

The taxonomic redefinition of *Clibanornis* (Claramunt et al. 2013) and the separation of *C. rufipectus* (Krabbe 2008) left unresolved the hidden diversity within the only polytypic species of the genus, *C. rubiginosus*. We addressed that gap with vocal evidence, and four species emerge from that analysis, supported to different degrees. *Clibanornis cinnamomeigula* is the most divergent member of the complex, with vocal, phenotypic, and genetic evidence converging on its recognition. *Clibanornis watkinsorum* is distinguished by its diagnostic introductory note and by its phylogenetic position apart from the remaining Amazonian taxa. *Clibanornis rubiginosus sensu stricto* and *C. obscurus* are separated by a discrete vocal character, and their ranges lie on opposite sides of the Andes. Phylogenetic corroboration and sampling in contact zones will refine these limits, and *caquetae* remains unstudied and provisionally placed with *obscurus*, but the vocal evidence presented here is the most complete available to inform a new taxonomy of the group.

### Proposed taxonomy and classification sequence

We recognize four species within what has been treated as *Clibanornis rubiginosus*, and describe below the diagnosis, taxon circumscription, and distribution of each. Although vernacular names fall outside the International Code of Zoological Nomenclature, we propose them for consistency of usage. We also comment on subspecies whose validity we consider doubtful (e.g., *moderatus*). The eight taxa we retain within *C. rubiginosus* sensu stricto were not separable in any acoustic variable, and assessing their diagnosability will require morphometric and genomic data.

### *Clibanornis rubiginosus* sensu stricto (Sclater, 1857)

Proposed English name: Rusty Foliage-gleaner

Protonym: *Anabates rubiginosus*

Holotype: NHMUK 1889.5.20.328

Type locality: “Cordova” (= Córdoba), Veracruz, Mexico.

Diagnosis: Song of a single note, lacking the introductory note present in every population east of the Andes. Total duration 0.43 ± 0.13 s and peak frequency 2087 ± 187 Hz, shorter and higher pitched than in *C. obscurus* and *C. watkinsorum*, and without the serrated vibrato of *C. cinnamomeigula*. Upperparts brownish with a darker crown, rufous tones on rump and tail, throat paler than the breast, belly olive-brown.

Remarks: We restrict the name to the clade west of the Andes, diagnosed by a single-note song. Variation within the species is expressed mainly in the intensity of rufous on the head, face, upperparts, throat, and breast, and separates two phenotypic subgroups. The Central American subspecies form a coherent if not homogeneous group, whereas the west of Andes South American subspecies *nigricauda*, *saturatus*, and *sasaimae* are darker or more heavily marked than any others in the complex and were formerly treated as a separate species (Chapman 1917, 1926, Cory and Hellmayr 1925) before being lumped by Peters (1951) and all subsequent authors. In addition, *umbrinus* is probably not reliably distinguishable from *veraepacis* (Remsen 2003). Songs of *fumosus* sit nearer the South American subspecies than the Central American ones in our ordination, a pattern that conflicts with its plumage similarity to *veraepacis* (Claramunt et al. 2013). Molecular data are consistent with that earlier treatment: *nigricauda*, *saturatus*, and *sasaimae* form a clade (Claramunt et al. 2013), and *nigricauda* falls apart from Central American *veraepacis* (Harvey et al. 2020). We retain all eight subspecies because our vocal data separate none of them, although the South American subspecies of this group are candidate for future recognition.

Distribution: Southern Mexico south through Central America to northwestern South America, entirely west of the Eastern Cordillera. Mostly premontane forest (including pine-oak forests in Mexico), extending south through the inter-Andean valleys, and to the foothills and lowlands of the Chocó in western Ecuador.

Polytypic, with eight subspecies: *rubiginosus* (Sclater, 1857), *guerrerensis* (Salvin and Godman, 1891), *veraepacis* (Salvin and Godman, 1891), *umbrinus* (Salvin and Godman, 1891), *fumosus* (Salvin and Godman, 1891), *nigricauda* (Hartert, 1898), *saturatus* (Chapman, 1915), and *sasaimae* (Meyer de Schauensee, 1947). No two were separable in any acoustic variable, and we propose retaining all eight pending modern phenotypic assessment and phylogeographic analysis.

English name: As redefined, *C. rubiginosus* no longer includes the populations east of the Andes, and reusing an established name for a smaller entity propagates confusion. Rusty Foliage-gleaner retains the color-based character of the current name and reflects the range of rusty to reddish plumage tones across the taxa west of the Eastern Cordillera.

### Clibanornis cinnamomeigula (Hellmayr, 1905)

Proposed English name: Piedemonte Llanero Foliage-gleaner

Protonym: *Automolus cinnamomeigula*

Holotype: AMNH 524305

Type locality: “Bogotá”, uncertain.

Diagnosis: Song of two subunits, the second modulated as five to seven rapid serrated oscillations heard as a resonant vibrato, a structure found in no other taxon in the genus. Peak frequency 2139 ± 243 Hz, the highest in the complex. Introductory subunit brief and dry with a clicking quality (0.07 ± 0.02 s), shorter than the introductory note of any population of *C. obscurus* or *C. watkinsorum*. Plumage uniformly cinnamon-rufous below except for a buff-rufous belly center; irides whitish to pale buff yellow against a dark face, a character unique within the complex.

Remarks: Hellmayr described *cinnamomeigula* as a species from a Bogotá trade skin without a precise locality or soft-part data, based on consistent differences in plumage pattern and intensity and in body proportions, with a longer and heavier bill. Chapman (1917, 1926) retained it at species rank, but Cory and Hellmayr (1925) treated it as a race of *rubiginosus*, and Hellmayr himself later wrote that this form and the two following ones were merely subspecies of the *rubiginosus* group. Molecular data place *cinnamomeigula* among the taxa east of the Andes in both studies available (Claramunt et al. 2013, Harvey et al. 2020), and never with the trans-Andean populations as implied by Cory and Hellmayr (1925). Its distinctively bright cinnamon rufous overall plumage alone sets it apart; and the characters that make it so distinctive, namely the saturation of the plumage and the contrast of the pale iris against the dark face, were documented in the field only from 2005 onward (pers. obs.). Had they been known in the nineteenth or early twentieth century, this taxon would hardly have passed as just one more geographic variant of the group.

Distribution: Eastern flank of the Eastern Cordillera of Colombia, along the foothills above the Llanos in extreme southeastern Cundinamarca, western Meta, southern Boyacá, and western Casanare; not yet recorded from Arauca but possibly extending there. Monotypic.

English name: Foliage-gleaner refers to the specific region the bird is endemic to, the transition between the eastern Llanos of the Orinoco basin and the eastern slope of the Eastern Cordillera. The *piedemonte llanero* of Colombian usage, referring to the Llanos foothills, is a proper name of easy referral.

### Clibanornis watkinsorum (Hellmayr, 1912)

Proposed English name: Peruvian Foliage-gleaner (but see Piacentini et al. 2015 for Watkins’s Foliage-gleaner)

Protonym: *Automolus watkinsi* Hellmayr, 1912

Holotype: ZSM 12628, Zoologische Staatssammlung München Type locality: Yahuarmayo, Marcapata, Peru.

Diagnosis: Song with the lowest peak frequency (1838 ± 132 Hz) and the longest total duration (0.66 ± 0.05 s) in the complex. Introductory note longer (0.21 ± 0.02 s) and lower pitched (1621 ± 162 Hz) than in the populations of *C. obscurus* we could sample, and that note separates it from *brunnescens* and *obscurus* in a discriminant function analysis. Little rufous overall, with dark scaling on the throat and a rufous nape; crown dark reddish chestnut; chin and upper throat ochraceous and conspicuously paler than the lower throat and malar area; belly medium olive-brown, flanks without a rufescent tinge (Remsen 2003, Claramunt et al. 2013).

Remarks; Hellmayr (1912) described this taxon as a species based on characters he considered sufficiently marked to separate it from the other Amazonian subspecies then known, and named it after its two discoverers, the brothers Henry and Casimir Watkins; the original spelling *watkinsi* is an incorrect original spelling (Costa 2017). Hellmayr (1925) later reconsidered, treating *watkinsi* and *cinnamomeigula* as races of the *rubiginosus* group. The evidence presented here reverses that conclusion, and species rank had already been restored by the Brazilian ornithological records committee (Piacentini et al. 2015).

Synonym: *Automolus moderatus* Zimmer, 1935, junior synonym. Zimmer described *moderatus* from a single specimen from Moyobamba, San Martín, northern Peru, and interpreted it as morphologically intermediate among several taxa of the complex. Later authors treated it as part of a variable *watkinsi* (Remsen 2003). The one recording we could find of alleged *moderatus* falls with *watkinsorum* in both introductory note duration and peak frequency.

Distribution: Western Amazonian lowlands and foothills, south of the Amazon and Marañon rivers, through much of Peru south to northern Bolivia and east into Acre, Brazil (Remsen 2003). Monotypic.

English name: Peruvian Foliage-gleaner anchors the species to the Andean foothills and western Amazonia of Peru, although its range extends into western Brazil and northwestern Bolivia.

### Clibanornis obscurus (Pelzeln, 1859)

Proposed English name: Guianan Foliage-gleaner (but see Piacentini et al. 2015 for Dusky Foliage-gleaner)

Protonym: *Anabates obscurus* Pelzeln, 1859

Syntypes

Naturhistorisches Museum Wien (catalogue number not traced). Type locality: Cayenne (purchased), French Guiana.

Diagnosis: Song of two notes, a short introductory note followed by a longer main note, which separates every population of this species from all of *C. rubiginosus sensu stricto*. Peak frequency higher and introductory note shorter than in *C. watkinsorum*; no serrated vibrato, which separates it from *C. cinnamomeigula*. Plumage dark rufescent brown above, throat and upper breast dark rufous, belly rufescent olive-brown.

Remarks: The name *obscurus* (1859) is the oldest available in this assemblage, ahead of *brunnescens* (1927), *venezuelanus* (1947), and *caquetae* (1947). The four taxa differ mainly in the intensity of rufous dorsally and toward the throat and malar region and in body size, with *obscurus* at the smallest extreme, but no diagnostic differences separate them (Claramunt et al. 2013, contra Meyer de Schauensee 1947). Of the four taxa we place here, only *obscurus* has been sequenced in a genome-scale dataset, where it falls with *cinnamomeigula* east of the Andes (Harvey et al. 2020). Claramunt et al. (2013) recovered their Guianan and northwestern Amazonian samples as close relatives. This agrees with their indistinguishable songs we report, so the vocally coherent unit is the entire assemblage rather than any subspecies within it. Species rank for *obscurus* is already in use for the Brazilian avifauna, with *venezuelanus* as a subspecies (Piacentini et al. 2015), and our results extend that circumscription to *brunnescens* and, provisionally, *caquetae*, for which no confirmed recording exists.

Distribution: Humid lowland and foothill forest of the Guianan Shield, southern Venezuela and northern Brazil, and western and central Amazonia from Colombia and Ecuador south, to possibly northern Peru, north of the Amazon river.

Polytypic, with four subspecies: *obscurus* (Pelzeln, 1859), *brunnescens* (Berlioz, 1927), *venezuelanus* (Zimmer and Phelps, 1947), and, provisionally, *caquetae* (Meyer de Schauensee, 1947).

English name: Guianan Foliage-gleaner points to the region encompassing the type locality and the core of the range, which extends across the lowlands of northern and central Amazonia to the Andean foothills in the west, north of the Amazon river.

## Acknowledgements

We thank Gary Stiles, Juan P. López-O., Óscar Laverde-R., and Santiago Claramunt for discussions at various stages of this work, and J. V. Remsen in particular for comments on an earlier draft. J. Camilo Váquiro, Natalia Pérez, Jonathan Espitia, Katherine Certuche, and especially Luis F. Recalde from the ORNIS research group, supported the study throughout. This work used the material archived by the Macaulay Library, Cornell Lab of Ornithology, and by xeno-canto, and we thank the recordists whose hard work we analyzed, each of whom is listed in Table S1. We thank the photographers listed in the Supporting information for the images in Fig. 2. We are grateful to the communities of Puerto Soya and Santa María, Boyacá (Catherine Acosta, Adrián Pinzón, and Eibar Algarra), and of Medina, Cundinamarca (Fredy Parra and Javier Beltrán), for their help during fieldwork and for their conservation effort along the Piedemonte Llanero. For access to type specimens, we thank the Academy of Natural Sciences of Philadelphia (J. Weckstein and N. Rice) and the American Museum of Natural History (B. T Smith and P. Sweet). The Instituto de Ciencias Naturales, Universidad Nacional de Colombia, provided institutional support.

