## Supplementary Material 1 for "Song divergence and a gleaming white iris reveal four species in a widespread Neotropical understory bird (*Clibanornis rubiginosus*, Furnariidae)"

**Appendix S1**. Source data for the photographs in Fig. 2, each with taxon, author, and archive accession where available. (a) C. r. rubiginosus, Alberto Lobato (ML205070211); (b) C. r. guerrerensis, Rick Bowers (ML622181692); (c) C. r. veraepacis, Daniel Garza T. (ML620037309); (d) C. r. umbrinus, José de León (ML325905941); (e) C. r. fumosus, Andrés Paniagua (ML625200058); (f) C. r. nigricauda, Lina Peña (ML632061571); (g) C. r. saturatus, Juan Carlos Herrera (ML610926164); (h) C. r. sasaimae, Andrés M. Cuervo (ML41086181); (i) C. r. caquetae, Andrés M. Cuervo, photograph of the holotype (ANSP 151330); (j) C. r. cinnamomeigula, Juan P. López-O., Santa María, Boyacá, Colombia, not archived; (k) C. r. brunnescens, Nick Athanas (ML164997171); (l) C. r. moderatus, Tomaz Melo (iNaturalist 38096364); (m) C. r. watkinsorum, J. Tobias (ML204088921); (n) C. r. obscurus, J. Mittermeier (Flickr 3747811332); (o) C. r. venezuelanus, Thiago Laranjeiras (WA2402259); (p) C. rufipectus, Johnnier Arango (ML610395523); (q) C. erythrocephalus, Stephan Lorenz (ML418950721); (r) C. rectirostris, Luciano Bernardes (ML611529893); (s) C. dendrocolaptoides, Martjan Lammertink (ML259294161).
